# *Spodoptera exigua* caterpillar feeding induces rapid defense responses in maize leaves

**DOI:** 10.1101/108076

**Authors:** Vered Tzin, Yuko Hojo, Susan R. Strickler, Lee Julia Bartsch, Cairo M. Archer, Kevin R. Ahern, Shawn A. Christensen, Ivan Galis, Lukas A. Mueller, Georg Jander

## Abstract

Insects such as beet armyworm caterpillars (*Spodoptera exigua*) cause extensive damage to maize (*Zea mays*) by consuming foliar tissue. Maize plants respond to such insect attack by triggering defense mechanisms that involve large changes in gene expression and the biosynthesis of specialized metabolites and defense signaling molecules. To investigate dynamic maize responses to herbivore feeding, leaves of maize inbred line B73 were infested with *S. exigua* caterpillars for 1 to 24 hours, followed by comprehensive transcriptomic and metabolomic characterization. Our results show that the most significant gene expression responses of maize to *S. exigua* feeding occur at early time points, within 4 to 6 hours after caterpillar infestation. However, both gene expression and metabolite profiles continued changing during the entire 24-hour experiment while photosynthesis genes were gradually decreased. The primary and specilaze metabolism shift maught be temporal and dynamic processes in the infested leaf tissue. We analyzed the effects of mutating genes in two major defense-related pathways, benzoxazinoids (*Bx1* and *Bx2*) and jasmonic acid (*Lox8*), using *Dissociation* (*Ds*) transposon insertions in maize inbred line W22. Together, these results show that maize leaves shift to implementation of chemical defenses within one hour after the initiation of caterpillar attack. Thus, the induced biosynthesis of specialized metabolites can have major effects in maize-caterpillar interactions.

**HIGHLIGHT:** A comprehensive transcriptic and metabolomic profiling time course of maize foliar responses to caterpillar feeding identifies genes for the synthesis of benzoxazinoids and phytohormones.

## INTRODUCTION

Plants perceive herbivory through mechanical cues from feeding damage, oviposition, and even insects walking on the leaf surface [1, 2], as well as through chemical cues from insect oral secretions and frass [3, 4]. In response to insect attack, up-regulated plant signaling pathways lead to the production of defensive metabolites. Two major signaling pathways that modulate plant defense against herbivores are regulated by salicylic acid (SA) and jasmonic acid (JA) derivatives[5, 6]. Jasmonates, in particular, regulate the production of toxic metabolites and a wide variety of other responses to insect herbivory [7-9].

In some graminaceous plants, including maize (*Zea mays*), wheat (*Triticum aestivum*), and rye (*Secale cereale*), JA induces the production of benzoxazinoids, a class of metabolites that can provide protection against insect herbivores, pathogens, and competing plants [10-13]. In maize, a series of nine enzymes (Bx1-Bx9) catalyze the biosynthesis of 2,4-dihydroxy-7-methoxy-1,4-benzoxazin-3-one glucoside (DIMBOA-Glc) from indole-3-glycerol phosphate [14, 15]. A family of three *O-*methyltransferases (Bx10-Bx12) methylate DIMBOA-Glc to form 2-hydroxy-4,7-dimethoxy-1,4-benzoxazin-3-one glucoside (HDMBOA-Glc) [16]. DIMBOA-Glc and HDMBOA-Glc are the most prevalent benzoxazinoids in maize seedlings [12, 14], though their relative abundance is quite variable among different maize inbred lines [16]. Recently, two additional enzymatic steps in this pathway were identified: a 2-oxoglutarate-dependent dioxygenase (Bx13) that catalyzes the conversion of DIMBOA-Glc into 2,4,7-trihydroxy-8-methoxy-1,4-benzoxazin-3-one glucoside (TRIMBOA-Glc) and an *O*-methyltransferase (Bx14) that converts 2,4-dihydroxy-7,8-dimethoxy-1,4-benzoxazin-3-one glucoside (DIM2BOA-Glc) into 2-hydroxy-4,7,8-trimethoxy-1,4-benzoxazin-3-one glucoside (HDM2BOA-Glc) [17]. Feeding by chewing herbivores brings benzoxazinoid glucosides into contact with β- glucosidases, leading to the formation of toxic breakdown products [15, 18, 19]. Induced benzoxazinoid accumulation and methylation of DIMBOA-Glc to produce HDMBOA-Glc in response to caterpillar feeding [10, 20] has been associated with increased resistance to several lepidopteran herbivores, including *Spodoptera exigua* (beet armyworm), *Spodoptera littoralis* (Egyptian cotton leafworm), *Spodoptera frugiperda* (fall armyworm), and *Diatraea grandiosella* (southwestern cornborer) [11, 15, 21, 22].

In the present study, we aimed to elucidate the dynamic of plant responses to *S. exigua* caterpillar feeding by integrating gene expression using high-throughput RNA sequencing with phytohormone and metabolite assays. Our study is focused on *S. exigua*, which has a wide host range, occurring as a serious pest of grains, vegetables, flower crops, and occasionally trees [23]. Maize inbred line B73 leaves were infested with *S. exigua* caterpillars for the time periods up to 24 hours, and statistical approaches were used to identify patterns of gene expression and metabolite changes. We confirmed the function of the specific benzoxazinoid and JA biosynthesis genes in maize-caterpillar interactions with *Ds* transposon insertions in inbred line W22.

## MATERIALS AND METHODS

### Plants and growth conditions

Single maize seeds were planted a 7.6 x 7.6-cm plastic pots (200 cm^3^), 1.5 cm deep, filled with moistened maize mix soil [produced by combining 0.16 m^3^ Metro-Mix 360, 0.45 kg finely ground lime, 0.45 kg Peters Unimix (Griffin Greenhouse Supplies, Auburn NY, U.S.A, 68 kg Turface MVP (Banfield-Baker Crop., Horseheads NY, USA), 23 kg coarse quartz sand, and 0.018 m^3^ pasteurized field soil]. Plants were grown for two weeks in growth chambers under a controlled photoperiod regime with a 16-h-light/8-h-dark cycle, 180 mMol photons/m^2^/s light intensity at constant 23 °C and 60% humidity.

### Caterpillar bioassays

*Spodoptera exigua* eggs were purchased from Benzon Research (Carlisle, PA, USA). After incubation for 48 h in a 29 °C incubator, first instar caterpillars were transferred to artificial diet (Beet Armyworm Diet, Southland Products Inc., Lake Village, AR, USA). Control and experimental maize seedlings received clip cages on the third leaf for 24 hours. For measuring the effect of caterpillar feeding on the maize transcriptome and metabolome, 2^nd^ to 3^rd^ instar *S. exigua* caterpillars were added to the clip cages for the final 1, 4, 6, or 24 hr of the experiment. All plant material was harvested at the same time. For *bx1::Ds* and *bx2::Ds* maize seeding caterpillar bioassays, individual caterpillars were confined on 10-day-old plants with micro-perforated polypropylene bags (15 cm × 61 cm; PJP Marketplace, http://www.pjpmarketplace.com) and caterpillar fresh weight was measured four days after the start of infestation. For *Lox8* (*ts1*) knockout lines, 3^rd^ instar caterpillars were placed in clip cages on the distal part of the 3^rd^ seedling leaf for 24 hr (*lox8(ts1)::Ds*) or ten days (*lox8/ts1-ref*), and caterpillar fresh weight was measured.

### Total RNA extraction

Leaf material was harvested, flash-frozen in liquid nitrogen, and ground to a fine powder using a paint shaker (Harbil, Wheeling, IL, USA) and 3 mm steel balls. The samples were homogenized, RNA was extracted using TRI Reagent (Sigma, St. Louis) and purified with the SV Total RNA isolation kit with on-column DNase treatment (Promega, Madison, WI, USA). Total RNA concentration and quality were assessed using a NanoDrop instrument (2000c; Thermo Fisher Scientific Inc. Waltham, MA USA).

### Transcriptome Sequencing, RNAseq data analysis and qRT-PCR analysis

Tissue from three individual maize plants was combined into one experimental replicate, and four replicates were collected for each time point. The purified total RNA (2-3 μg) was used for the preparation of strand-specific RNAseq libraries [24, 25] and amplified for 16 cycles. The purified RNAseq libraries were quantified, and 20 ng of each was used for next generation sequencing using an Illumina HiSeq2000 instrument (Illumina, San Diego, CA) at the Weill Medicine School Facility (Cornell University, NY, USA) with a 101 bp single-end read length. Libraries were multiplexed and sequenced in one lane. Read quality values were checked using FastQC (http://www.bioinformatics.babraha-m.ac.uk/projects/fastqc). Low-quality sequences and adapters were trimmed and removed using Fastq-mcf (http://ea-utils.googlecode.com/svn/wiki/FastqMcf.wiki), with a minimum length of 50 bp and minimum quality value of 30. RNAseq analysis was performed following the protocol published by Anders et al. 2013 [26], using the maize genome version B73 AGP v3.22 as a reference [25]. The benzoxazinoid genes were also analyzed using AGP v3.20 to determine the values of *Bx7* and *Bx13*, which were excluded from the AGP v3.22 as low-confidence genes. Reads were mapped with the Tophat2 [27] and Cuffdiff packages [28] of Cufflinks version 2.2.1, using the geometric mean option. Transcripts showing at least one FPKM (Fragment per Kilobase of exon per Million fragments)of transcript in 3 or more replicates for each time point were kept for differentially expressed gene detection.

To verify the results of the RNA-seq analysis, an additional experiment was conducted, and total RNA was extracted. First strand cDNA was synthesized by M-MLV reverse transcriptase (TaKaRa) with a new biological experiment of RNA samples. qRT-PCR was conducted as described previously [29], and the primer list of the seven genes is described in Supplementary Table S1.

### Targeted and untargeted metabolite assays

For assays of maize metabolites, approximately 2 cm of leaf material on which caterpillars had been feeding was collected from the third leaf, as well as from control plants without caterpillars. For non-targeted metabolite assays, frozen powder of fresh tissue was weighed in a 1.5-ml microcentrifuge tube, and extraction solvent (methanol/water/formic acid, 70:29.9:0.1, v/v) 1:3 ratio was added [30]. The tubes were vortexed briefly, shaken for 40 min at 4 ºC, and centrifuged for 5 min at 14,000 *g.* The samples were filtered through a 0.45-micron filter plate (EMD Millipore Corporation) by centrifuging at 2,000 *g* for 3 min and the supernatant was diluted 1:9, with the extraction solvent then transferred to an HPLC vial. LC-MS/MS analysis was performed on a Dionex UltiMate 3000 Rapid Separation LC System attached to a 3000 Ultimate diode array detector and a Thermo Q Exactive mass spectrometer (Thermo Scientific). The samples were separated on a Titan C18 7.5 cm x 2.1 mm x 1,9 μm Supelco Analytical Column (Sigma Aldrich), as previously described [17]. For data analysis, raw mass spectrometry data files were converted using the XCMS [31] followed by CAMERA R package [32]. The chromatographic peaks were compared with the retention time, accurate mass and UV spectrum of standards of DIMBOA, DIMBOA-glucoside and HDMBOA-glucoside. Other benzoxazinoids were identified based on their accurate masses and UV spectra.

### Phytohormone analysis

Maize leaves (30-100 mg fresh weight) were harvested, frozen in liquid nitrogen, and lyophilized. Samples were homogenized in a FastPrep®-24 (MP Biochemicals, Santa Ana, CA) using five 2.3 mm zirconia beads and 1 ml ethyl acetate solvent spiked with deuterated internal standards (IS) (25 ng d3-JA, 5 ng d3-JA-Ile, 10 ng d6-ABA, and 20 ng d4-SA). Samples were centrifuged for 15 min at 13,200 *g*, 4ºC, and supernatants were collected in clean 2 ml microcentrifuge tubes. Pellet extraction was repeated once with 0.5 mL pure ethyl acetate and vortexing for 5 min at 23 °C, and supernatants were pooled with the previous fraction after centrifugation, as before. Samples were dried completely under vacuum in a miVac Quatro concentrator (Genevac Ltd, Ipswich, UK). Each sample was dissolved in 300 µL 70% methanol/water (v/v) and vortexed for 5 min at 23 °C. Then, 1,700 µl buffer (84 mM ammonium acetate; pH 4.8) was added to each sample prior to application and retention of phytohormones on preconditioned 3 ml SPE columns (Bond Elut-C18, 200 mg, Agilent Technologies Inc., Santa Clara, CA, USA) set in a QIAvac 24 Plus system (QIAGEN, Germantown, MD, USA). After brief drying with an air stream, samples were eluted with 800 µl 85% methanol/water (v/v) into clean 1.5 ml microcentrifuge tubes. After brief spin at 12,000 *g* to remove insoluble materials,10 µL aliquots were analysed using a triple quadrupole LC-MS/MS 6410 system (Agilent Technologies) equipped with a Zorbax SB-C18 column [2.1 mm id × 50 mm, (1.8 µm), Agilent Technologies] kept in thermostat-controlled chamber at 35°C. The solvent gradient, A (0.1% formic acid in water) vs. B (0.1% formic acid in acetonitrile), was used as follows: 0 min,15% B; 4.5 min, 98% B; 12 min 98% B; 12.1 min, 15% B; and 18 min,15% B, at a constant flow rate 0.4 mL/min. Mass transitions, hormone/ Q1 precursor ion (m/z)/ Q3 product ion (m/z), were monitored for each compound as follows: JA/209/59, JA-Ile/322/130, abscisic acid (ABA)/263/153, SA/137/93, 12-oxophytodienoate (OPDA)/291/165, OH-JA/225/59, OH-JA-Ile/338/130, COOH-JA-Ile/352/130, JA-Val/308/116, d3-JA/212/59, d3-JA-Ile/325/130, d6-ABA/269/159, and d4-SA/141/97. The fragmentor (V)/ collision energy (V) parameters were set to 100/6 for JA, OH-JA, and OPDA; 135/15 for JA-Ile, OH-JA-Ile, COOH-JA-Ile and JA-Val; 130/5 for ABA; and 90/12 for SA. JA, JA-Ile, ABA and SA amounts were directly calculated from the ratio of the endogenous hormone peak and the known deuterated internal standard. Compounds for which the authentic deuterated standards were not available were quantified using their structurally nearest deuterated internal standard, and expressed as equivalents of this compound (OPDA as d3-JA eq.; hydroxy (OH)-JA-Ile, carboxy (COOH)-JA-Ile, and JA-Val as d3-JA-Ile eq.). Phytohormone concentrations were calculated relative to the actual fresh mass of each sample used for extraction.

### Isolation of transposon insertions knockout lines and genotyping

The *lox8/ts1-ref* mutant was acquired from the Maize Genetics Cooperation Stock Center at The University of Illinois at Urbana-Champaign (Maize COOP, http://maizecoop.cropsci.uiuc.edu) as a segregating 1:1 heterozygous: mutant population and selfed to generate a 1:2:1 segregating popuation, as describe previously [9]. The *Ds* transposon insertions in the W22 maize line and were identified in the following genes of interest through the *Ac/Ds* tagging project website (http://www.acdstagging.org) [33]. Seed stocks for *bx1::Ds* (Gene ID- GRMZM2G085381; Ds-B.W06.0775), *bx2::Ds* (Gene ID - GRMZM2G085661; Ds-I.S07.3472) and *lox8(ts1)::Ds* (Gene ID - GRMZM2G104843; Ds-B.S06.0143). The primer list and Maize Genetics Cooperation Stock Center information is in Supplementary Table S2.

### Statistical analysis

Partial Least Squares Discriminant Analysis (PLS-DA) was conducted as previously described [34] and the plot was drawn using MetaboAnalyst 3.0 software [35]. Venn diagrams were made using the Venny 2.1.0 drawing tool (http://bioinfogp.cnb.csic.es/tools/venny/index.html). The optimal number of clusters for the transcriptomic data for K-means clustering analysis was calculated using Gap [36] and NbClust R packages [37]. The *K-*mean analysis was performed on scaled and centered FPKM Log2 values and presented in standard (Z) score format. Gene ontology enrichment analysis was conducted using the PlantGSEA tool (http://structuralbiology.cau.edu.cn/PlantGSEA/) [38]. Fisher’s exact test was used to take into account the number of genes in the group query, the total number of genes in a gene set, and the number of overlapping genes, with the false discovery rate calculated using the Hochberg procedure (*P* value = 0.05). Statistical comparisons were made using JMP Pro 11 (SAS, www.jmp.com).

## RESULTS AND DISCUSSION

### Transcriptomic analysis of maize responses to caterpillar feeding

To investigate the global transcriptomic changes in response to caterpillar feeding, the third leaves of maize inbred line B73 seedlings were infested with two 2nd-3rd *S. exigua* instars for 0, 1, 4, 6, or 24 hr. A recent study of the effect of *S. littoralis* on maize defense mechanisms showed that induced herbivore resistance is highly localized and dependent on benzoxazinoid biosynthesis [39]. Therefore, we focused our transcriptomic assays on the caterpillar-infested section of the leaf. Caterpillar exposure was started in a staggered manner, such that all samples were harvested at the same time on the same day (Supplemental Fig. S1). Comparison of transcriptome data (Illumina RNAseq) to the B73 genomic sequence [25], which has predicted gene models (AGPv3.22; www.maizegdb.org), showed approximately 40,000 unique transcripts (Supplemental Table S3). The expression patterns of seven selected genes were confirmed by quantitative reverse transcriptase-PCR (qRT-PCR) using independently generated plant samples. Comparison of gene expression using these two methods showed a similar expression pattern and a high correlation coefficient (*R*-value; Supplemental Fig. S2). After data filtering (genes which had expression values of zero more than three times were excluded), approximately 20,000 transcripts from the RNAseq data set were analyzed for each of the four caterpillar-infested time points to detect genes that were differentially expressed relative to untreated control leaves (Supplemental Table S4). The gene expression levels were used to conduct a Partial Least Squares Discriminant Analysis (PLS-DA) for each of the biological replicates (Fig. 1A). Samples from each time point cluster with one another and the expression profiles separate gradually over time from the 0 hr (control) sample. Samples from the 24-hour time point clustered furthest from the control samples, indicating the greatest changes in gene expression after the onset of caterpillar feeding. Genes with significant expression differences (*P* value ≤ 0.05, FDR adjusted) and at least two-fold changes relative to the controls for at least one of the time points were selected for further analysis (Supplemental Table S5). After the initiation of caterpillar feeding, thousands of transcripts showed altered expression at each of the time points. The number of down-regulated genes increased gradually and, in the 24 hr sample, was similar to the number of the up-regulated genes (1,838 down-regulated and 1,954 up-regulated; Fig. 1B). The distribution of up- and down-regulated genes was calculated at each time point and present in the Venn diagram (Fig. 1C). Although a unique set of genes was increased at each time point (total 3,078), expression of a large number of genes (914) was induced in all time points. The comparison of genes altered by caterpillar feeding after 24 hr and 1 hr is presented in Supplemental Fig. S3.

Metabolic changes were studied in caterpillar-infested maize leaf samples using untargeted LC-MS/MS in negative (Supplemental Fig. S6) and positive (Supplemental Fig. S7) ion mode. PLS-DA clustering pattern of the untargeted metabolite analysis is presented in Figure 1D. The negative ion mode plot showed significant separation from the controls at 6 and 24 hr after initiation of caterpillar feeding, whereas the PLS-DA plot of the positive ion mode showed significant separation only after 24 hr. The overall similarity of the transcriptomic and metabolomic data suggested that caterpillar-induced gene expression changes lead to induced changes in the metabolome at the same or later time points.

**Figure 1.**
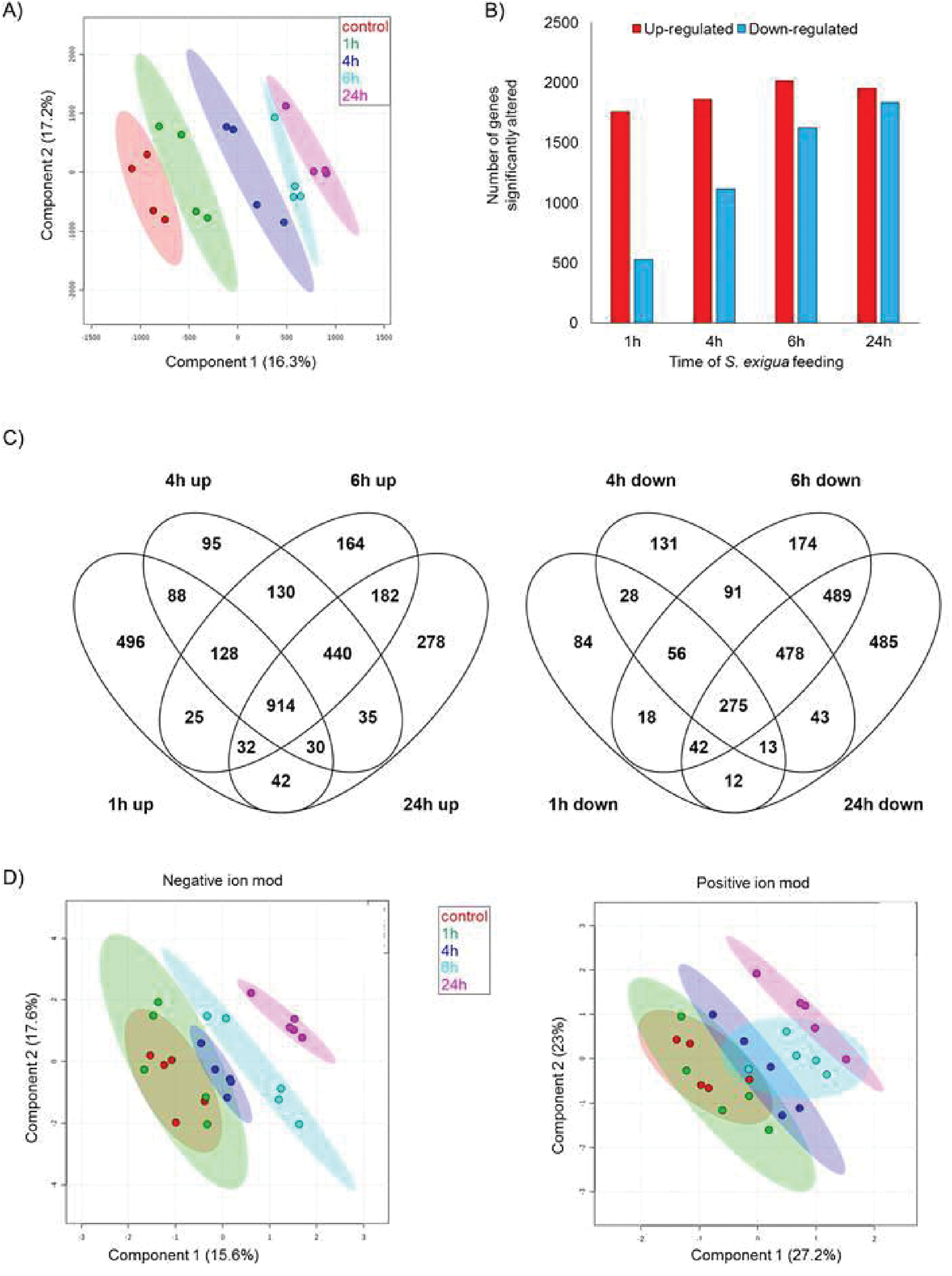
Transcriptomic and metabolomic overview of a time course of S. *exigua* feeding on maize inbred line B73 foliage. A) Partial least squares Discriminant Analysis (PLS-DA) generated from 20,825 genes (FPKM > 0 in at least 18 samples). Ovals indicate 95% confidence intervals. B) A total numbers of transcripts that were significantly up- or down-regulated; and C) Venn diagram illustrating the number of genes up- or down-regulated by Caterpillar infestation in the time course. *P* value < 0.05 FDR and fold change > 2 or < 0.5. D, E) Untargeted metabolomics of maize leaf responses to Caterpillar feeding. Partial least squares Discriminant Analysis (PLS-DA) plots identified by negative ion mode (2,044 mass Signals) and positive ion mode (1,917 mass Signals). Ovals indicate 95% confidence intervals.

### Clustering the transcriptome dataset

The significantly differentially expressed genes were subjected to *K*-Means clustering using Pearson Correlation distances (R). The *K*-Means analysis was performed on scaled and centered FPKM Log2 values, and each cluster is represented by the Z-score (standard score) of the gene expression of the set of genes showing similar response patterns to caterpillar herbivory. The 16 clusters were divided into four expression groups according to the trends of the standard score: i) clusters with strong increasing average (two standard deviations); ii) clusters with moderately increasing average (approximately one standard deviation), ii) clusters with moderately decreasing average (approximately one standard deviation), and iv) clusters with moderately decreasing average that significantly deviates from the population average (high FPKM) (Fig. 2). The distribution of genes into the 16 clusters is presented in Supplemental Table S8.

To elucidate the biological processes that contribute to each gene expression cluster, over-representation analysis was performed using PlantCyc output from PlantGSEA tool [38] (Table 1). The first pattern includes two clusters of genes (1 and 2) that are highly induced by caterpillar feeding. Although these clusters contain a relatively small number of genes (133 and 74 respectively), many pathways of transcripts associated with plant defense and stress responses were over-represented, including biosynthesis of phenylpropanoids, suberin, jasmonic acid, monosaccharides, methionine biosynthesis, and methionine degradation toward ethylene biosynthesis and *S*-adenosyl-*L*-methionine. The second pattern includes six clusters (3-8) of moderately increased gene expression, mainly associated with sucrose degradation and cellulose biosynthesis, as well as phenylpropanoids, suberin, JA, β-alanine, glutamine, TCA cycle, fatty acid biosynthesis, and cytokinin-O-glucoside biosynthesis. The moderately decreased gene expression pattern (clusters 9-14) contains genes associate with photorespiration, nitrogen fixation, and flavonoid biosynthesis. For the *K-*means analysis, we used the FPKM dataset. Therefore, we identified a pattern of genes with high FPKM and moderately decreased (pattern 4). These two clusters (15-16) include genes that are involved in the photosynthesis light reaction, Calvin-Benson-Bassham cycle, gluconeogenesis, and glycolysis (Table 1). However, this reduction in photosynthetic genes might be temporal follow by readjustment of the photosynthetic capacity under the biotic stresses [40]. Together, these observations indicate that there is a shift from primary metabolism toward the synthesis of defensive metabolites in response to *S. exigua* feeding on maize.

**Table 1.**
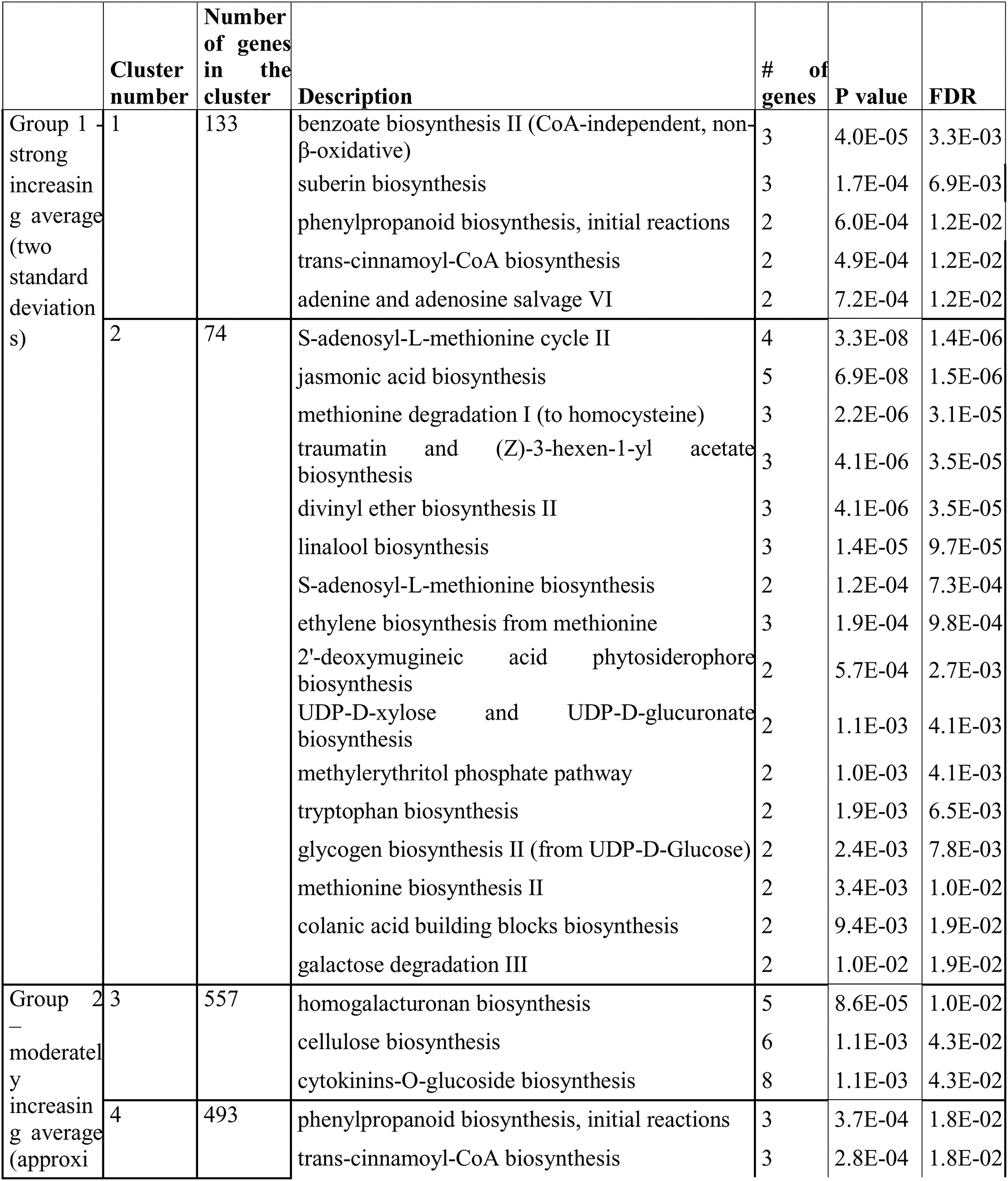
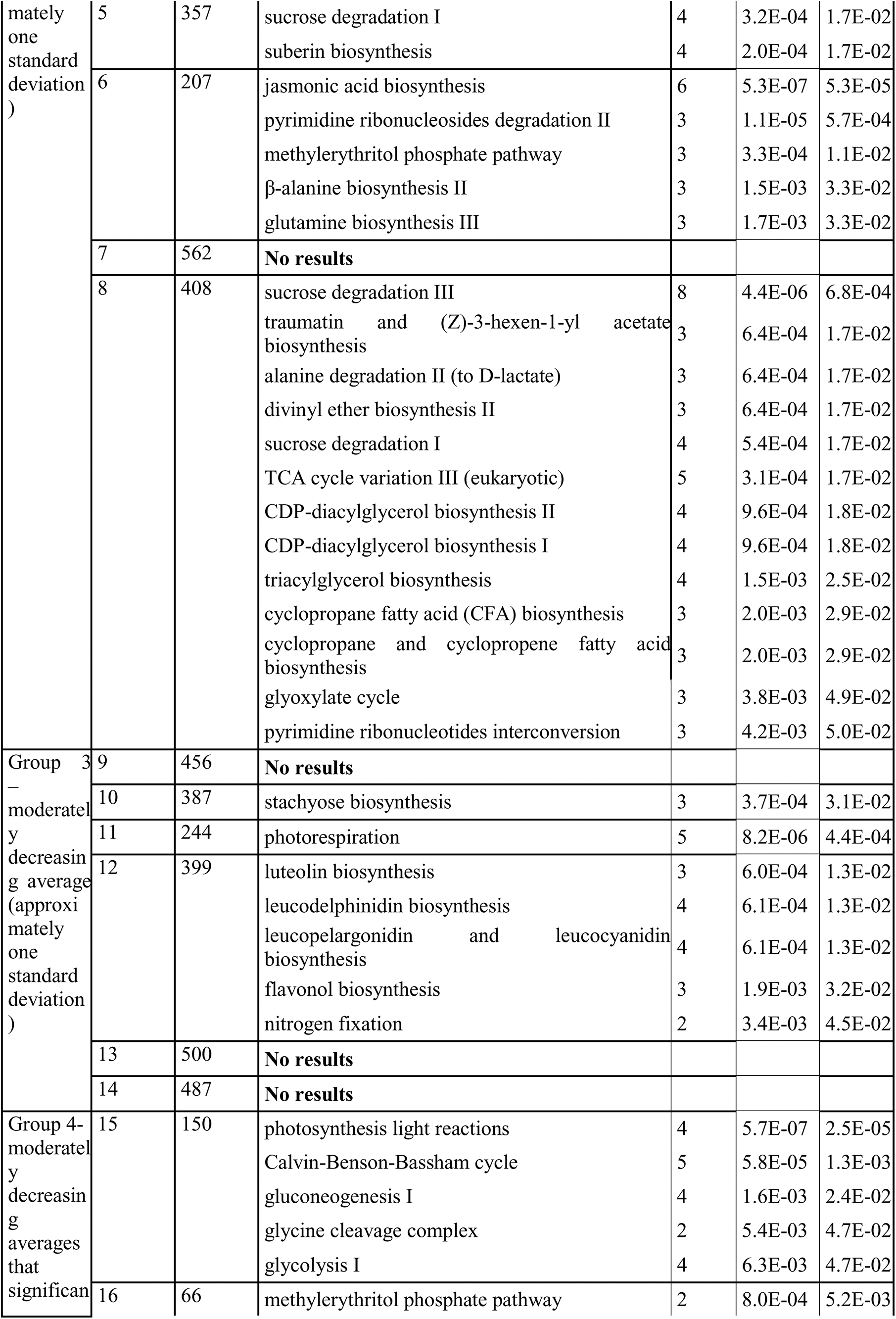
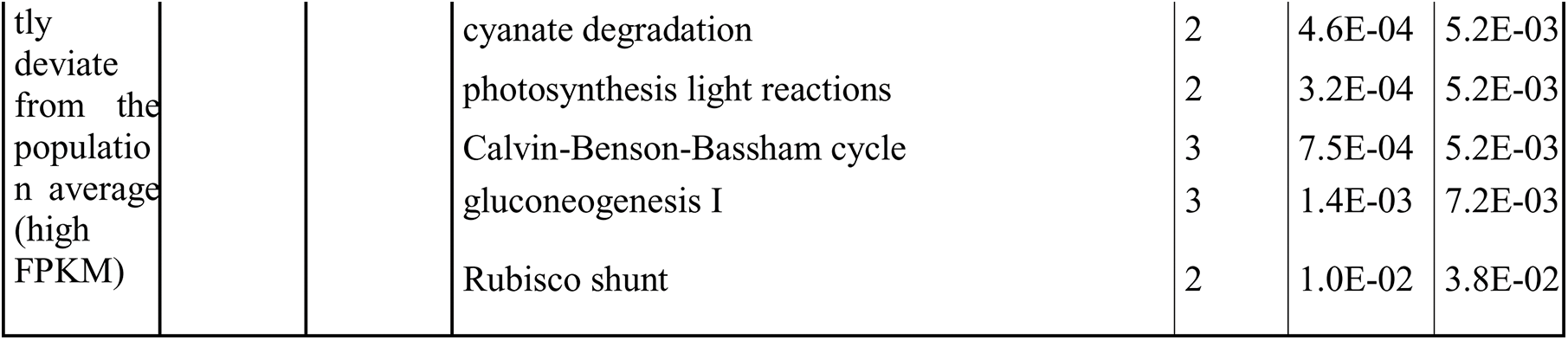
Enrichment analysis of metabolic pathways grouped by K-means clustering. Gene expression patterns were sorted into 16 clusters, as determined by *K*-Means analysis of transcripts detected in the B73 maize in inbred line at 0, 1, 4, 6 and 24 hr after caterpillar feeding.

**Figure 2.**
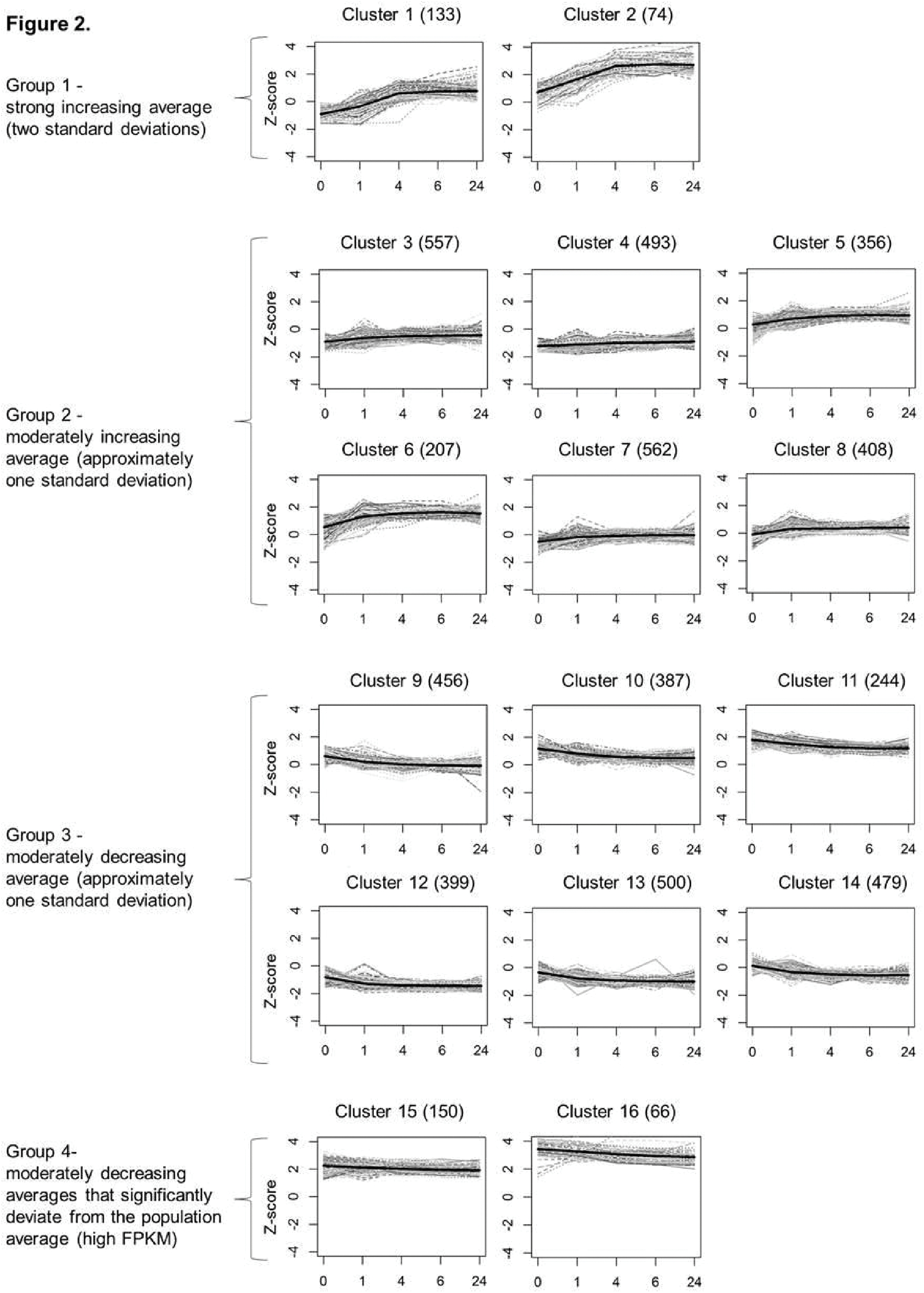
*K*-means clustering of genes expressed during caterpillar infestation. Gene expression in FPKM Log2 values after *S. exigua* feeding over a 0-24 h time course. Genes were selected according to the following parameters:*P* value < 0.05 FDR and fold change > +/-2 (5,470 genes). In bold: Z-score represents the number of standard deviations between the genes represented in the cluster and the mean of the population distribution. In brackets: the number of genes in each cluster.

### Plant hormone-related genes and metabolites induced by *Spodoptera exigua* damage

To identify the transcriptional signatures of hormonal responses in caterpillar-infested maize plants, the Hormonometer tool was used [41]. We evaluated similarities in the expression profiles elicited by caterpillar herbivory and those induced by application of the plant hormones methyl-jasmonate, 1-aminocyclopropane-1-carboxylic acid (a precursor for ethylene), ABA, indole-3-acetic acid (auxin), cytokinin (zeatin), brassinosteroid, gibberellic acid, and salicylic acid. As the hormone treatments were conducted with *Arabidopsis thaliana* (Arabidopsis), we selected the orthologous genes from Arabidopsis and the B73 genome. The genes included in the RNAseq analysis after the filtering processes contain a corresponding Arabidopsis Probeset ID, and a total of 10,242 Arabidopsis orthologs of maize genes were used as input for the Hormonometer analysis (Supplemental Table S8). As shown in Figure 3, genes associated with JA-, ABA-, auxin-, and SA-dependent signaling were highly induced after 1 hr of infestation, followed by moderate induction of these phytohormone signaling pathways at later time points (Fig. 3). Ethylene-, gibberellin-, cytokinin-, and brassinosteroid- responsive genes showed a negative correlation with caterpillar-induced genes and those that were induced within half an hour after hormonal treatment and moderately increased after 3 hr of hormonal treatment. A dendrogram analysis of the data showed that hormone-related gene expression changes gradually from 1 to 24 hr, and that responses in the first hour after caterpillar feeding are distinct from those observed at later time points. This suggests that major hormonal induction occurs within one hour after caterpillar infestation (Fig. 3).

We measured alteration in the phytohormones induced by *S. exigua* feeding using LC-MS/MS. As shown in Figure 4, the ABA level was significantly increased at 4 and 6 hr after the initiation of caterpillar feeding. Although SA levels showed a similar trend, the results were non-significant. The JA level was significantly increased from 1 to 24 hr, and other JA conjugates (JA-Val, JA-Ile, OH-JA-Ilu, and COOH-JA-Ile) were highly induced from 4- to 24 hr after caterpillar feeding. The JA precursor OPDA was only increased in abundance after 24 hr.

**Figure 3.**
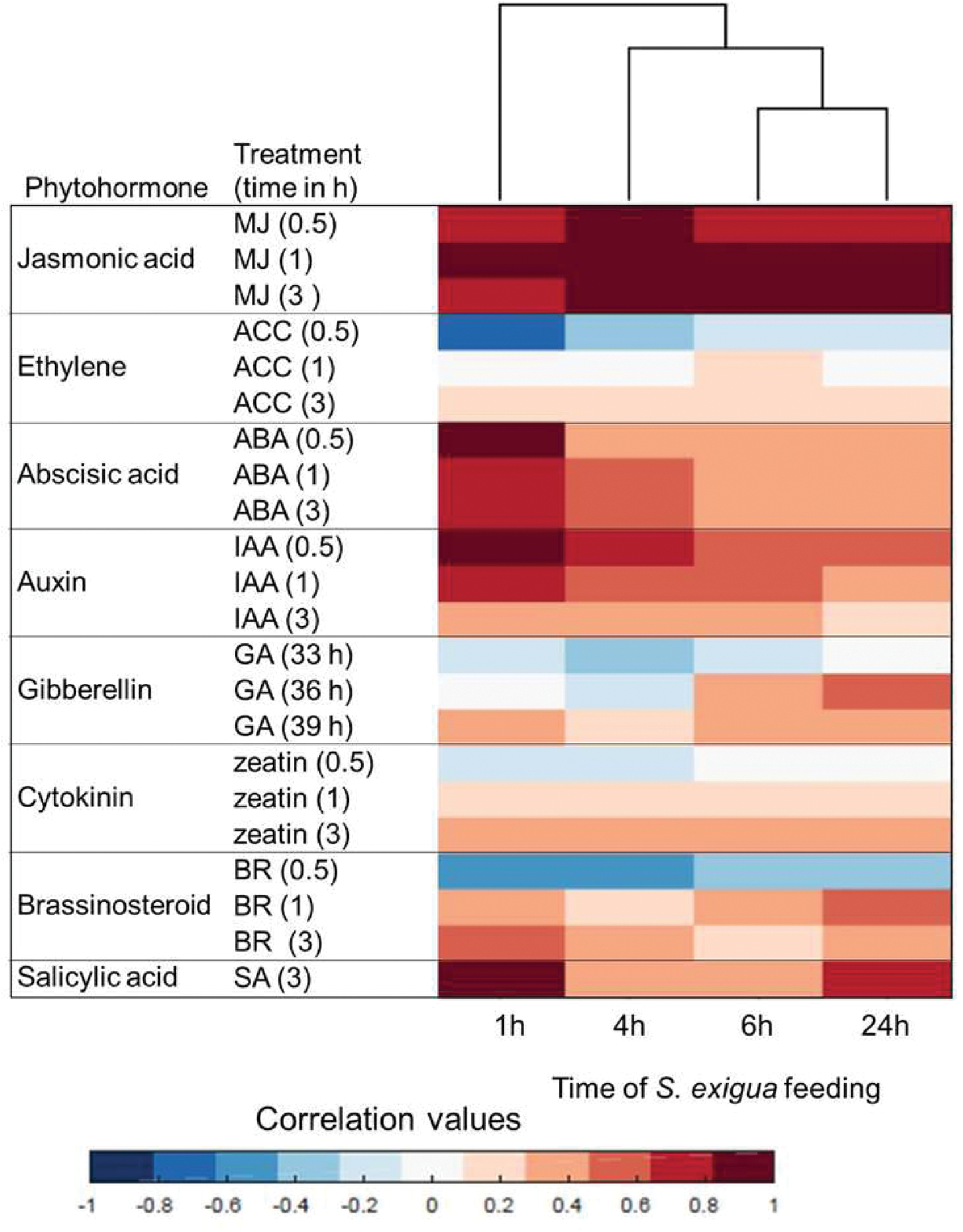
Plant hormone signatures based on transcriptomicdata generated after S. *exigua* feeding on maize leaves. Red, indicates a positive correlation between the maize *S. exigua* Caterpillar treatment and a particular hormone response; blue, negative correlation. MJ; methyl jasmonate, ACC; 1 -aminocyclopropane-1 -caroxylic acid (precursor of ethylene), ABA; abscisicacid, IAA; indole-3-aceticacid, GA; gibberelic acid, BR; brassinosteroid, SA; salicylic acid. The analysis was conducted using the Hormonometertool [41].

### Caterpillar-induced changes in jasmonic acid biosynthesis

Abundance of JA, SA and ABA was affected by *S. exigua* feeding (Fig. 4). However, the greatest induction at the gene expression and metabolite level was JA-related and occurred at 4 hr to the 24 hr after the start of infestation. Maize lipoxygenases (LOX) initiate fatty acid oxidation pathways for the synthesis of compounds that function in plant defense against insect herbivory [42, 43] (Supplemental Fig. S4A). Up-regulation of gene expression (Fig. 2), as well as hormonal response signatures (Fig. 3), and phytohormone quantification (Fig. 4) suggested that caterpillar feeding elicits production of a complex array of oxylipins. Therefore, we investigated the expression of genes associated with oxylipin and JA production [44, 45] http://www.plantcyc.org; Fig. 5A) in more detail. In general, the first steps of the pathway are highly induced by caterpillar feeding. Lipoxygenases that enable production of 12-oxo-phytodienoic acid (12-OPDA) and its downstream JA products (13-LOXs; *LOX7, 9, 10, 11* and *13*) were induced from 4 to 24 hr. In contrast, *LOX8/ts1* (GRMZM2G104843) was highly induced from the first hour of infestsion. A similar pathway involving 9-LOX activity on linolenic and linoleic acid (*LOX3*,*4*,*5* and *6*), which leads to the 12-OPDA positional isomer, 10-oxo-11-phytodienoic acid (10-OPDA) and 10-oxo-11-phytoenoic acid (10-OPEA) were highly induced. Allene oxide synthase (AOS) genes, which encode the second step of the jasmonic acid pathway, also were highly increased. In addition, allene-oxide cyclase (AOC) genes, which encode the third step in the pathway, were upregulated at all time points. Elevated JA levels have been associated with insect resistance in several plant species [9, 46, 47].

**Figure 4.**
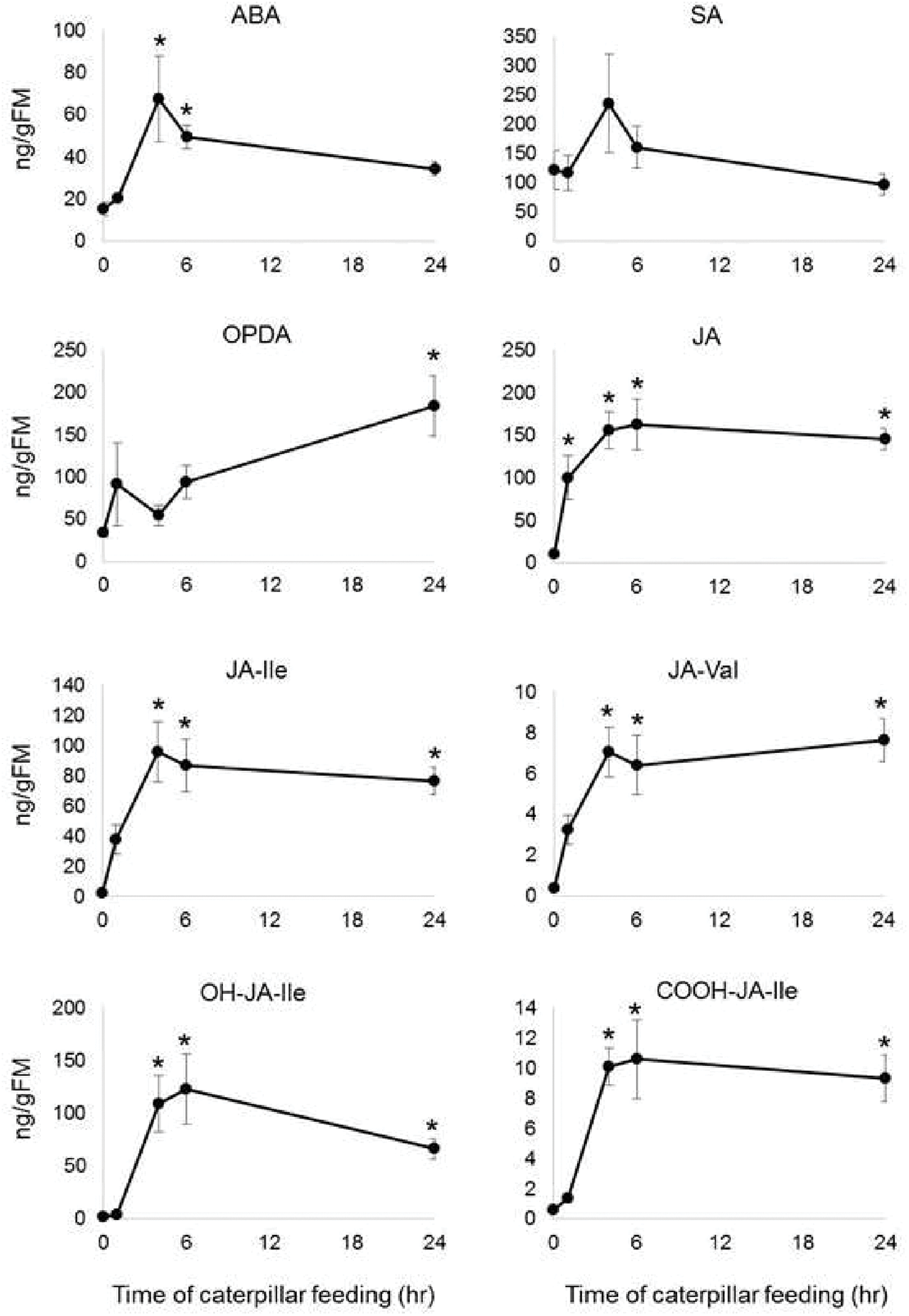
Plant phytohormones produced after *S. exigua* feeding on maize leaves. JA, jasmonicacid;ABA, abscisicacid; and SA, salicyiicacid. Mean +/- SE of n = 5. *P< 0.05 Student’s f-test relative to uninfested control.

The expression patterns of oxophytodienoate reductases (*OPR*), which encode the fourth step of the jasmonic acid pathway, were varied: *OPR7* was highly increased whereas *OPR1* and *OPR2* were slightly increased after caterpillar feeding. Moreover, the expression levels of *OPR3* and *OPR6* were reduced. The subsequent enzymatic steps, encoded by acyl-CoA oxidase, enoyl-CoA hydratase, acetyl-CoA *C*-acyltransferase and long-chain-3-hydroxyacyl-CoA dehydrogenase were slightly increased 4 hr and later after initiation of caterpillar feeding (Fig. 5A). This suggested that other intermediates of the oxylipin pathway also might have functions in plant defense.

With *LOX8* expression in response to *S. exigua* feeding predominating over the other maize 13-LOXs, we genetically investigated its role in caterpillar-induced jasmonate production using two different *LOX8/ts1* knockout alleles (*lox8/ts1::Ds* and *lox8/ts1-ref*). While several jasmonates were significantly less induced by *S. exigua* in *lox8/ts1::Ds* (Fig. 5B) and *lox8/ts1-ref* (Supplemental Fig. S4B), they were not completely absent, suggesting that multiple 13-LOXs can provide substrats induced jasmonate biosynthesis. These results are consistent with our expression data, which show that expression of multiple 13-LOXs is induced, and parallel previous findings in maize and Arabidopsis [9, 48]. Compared to wildtype controls, caterpillar weight gain was not significantly increased when feeding on *lox8* knockout lines (data not shown). Christensen et al (2013) showed increased in *S. exigua* bodyweight after feeding of *lox10* mutants, and suggested that although both LOX-8 and LOX-10 are 13-LOXs, they differ in their sub-cellular locations, providing substrate for the green leaf volatile and JA biosynthesis pathways, respectively [9].

Comparing the expression levels of oxylipin biosynthetic genes showed that 9-LOX genes are highly induced relative to 13-LOX genes (Fig. 5A). Whereas 13-LOX-derived linolenate oxidation produces 12-OPDA and JA, 9-LOX-derived linoleic acid oxidation enables the production of 10-oxo-11-phytoenoic acid 10-OPEA [44] and likely as yet unknown downstream products. In maize, fungal infection by southern leaf blight (*Cochliobolus heterostrophus*) causes induction of 9-LOXs and production of 10-OPEA, which displays local phytoalexin activity [49]. Therefore, we hypothesize that the 9-LOX branch of the oxylipin pathway also functions in direct defense or defense signaling in response to caterpillar feeding.

**Figure 5.**
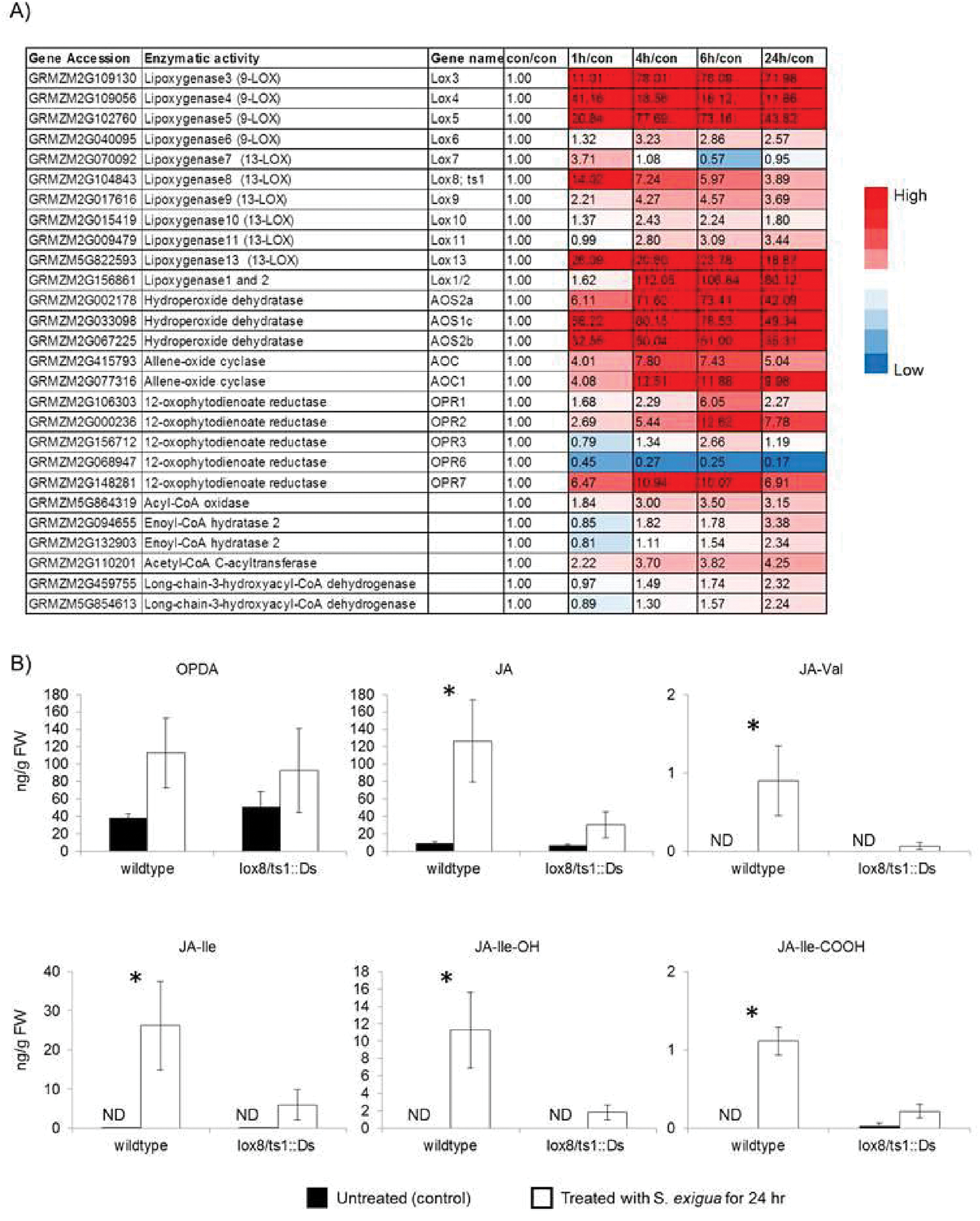
Effects of Caterpillar feeding on jasmonic acid biosynthesis. A) Heat map of gene expression that is related to jasmonic acid (JA) biosynthesis. Values are presented in fold change relative to untreated control. Mean +/- SE of n = 4. B) JA and JA conjugate levels of in a *lox8/ts1::Ds* gene knockout line in response to Caterpillar attack. Black bars, untreated, white bars- Caterpillar infestation for 24 hr. *P < 0.05, two-tailed Student’s *t-*test. ND = not detected.

### Benzoxazinoid biosynthesis is involved in herbivore defense mechanisms

The role of benzoxazinoids in defense against herbivory has been studied extensively in maize [12, 13, 50] (Fig. 6A). Therefore, we elaborated benzoxazinoid gene expression and function in response to *S. exigua* feeding (Fig. 6B and Supplemental Table S11). As shown in Figure 6B, transcripts of *Bx1*, *Bx*, *Bx3*, and *Bx6* were highly induced from 4 to 24 hr, whereas *Bx4*, *Bx5*, *Bx8*, *Bx9*, and *Bx7* were highly induced after 4 and 6 hr of caterpillar infestation. *Bx10, Bx11*, and *Bx13* were increased within one hour of caterpillar infestation, suggesting that an immediate response to caterpillar feeding is the conversion of DIMBOA-Glc to HDMBOA-Glc or DIM2BOA-Glc. However, *Bx14* was induced only after 4 and 6 hr of caterpillar infestation. Both DIMBOA-Glc and HDMBOA-Glc abundance gradually increased from 4 to 24 hr (Fig. 6C).

To further investigate the effect of *S. exigua* feeding on benzoxazinoid content, we employed the previously identified *bx1::Ds* and *bx2::Ds*, mutations in the W22 genetic background [29, 51]. DIMBOA-Glc and HDMBOA-Glc, were significantly increased in the W22 wildtype treated with caterpillars compare to untreated plants (Fig. 6D). However, the levels of these compounds were very low in the *bx1::Ds* and *bx2::Ds*, even with caterpillar infestation. At least two other maize genes, GRMZM2G046191 (*IGL1*), and GRMZM5G841619 (*TSA1*), encode the same indole-3-glycerol phosphate lyase enzymatic activity as *Bx1* [52, 53]. Thus, the absence of DIMBOA-Glc and HDMBOA-Glc induction by caterpillar feeding on the *bx1::Ds* mutant line indicates that there is either metabolic channeling or the other two genes are not strongly regulated in response to caterpillar feeding (Supplemental Table S5) The abundance of HDMBOA-Glc was significantly increased in *bx2::Ds* plants after caterpillar infestation. It is possible that the other cytochrome P450 enzymes in the pathway (Bx3, Bx4, or Bx5) catalyze the initial Bx2 indole oxidation reaction to a more limited extent.

Corn leaf aphids (*Rhopalosiphum maidis*) grow better on *bx1::Ds* and *bx2::Ds* mutant lines than on wildtype W22 [29, 51]. To determine whether this is also the case for *S. exigua*, caterpillar body weight was measured after four days of feeding on mutant and wildtype seedlings. There was a significant increase in caterpillar body mass on *bx1::Ds* and *bx2::Ds* mutant seedlings relative to wildtype W22 (Fig. 6E). A similar increase in body weight was observed with *S. littoralis* feeding on maize inbred line B73 *bx1* mutant plants relative to wildtype B73 [39].

**Figure 6.**
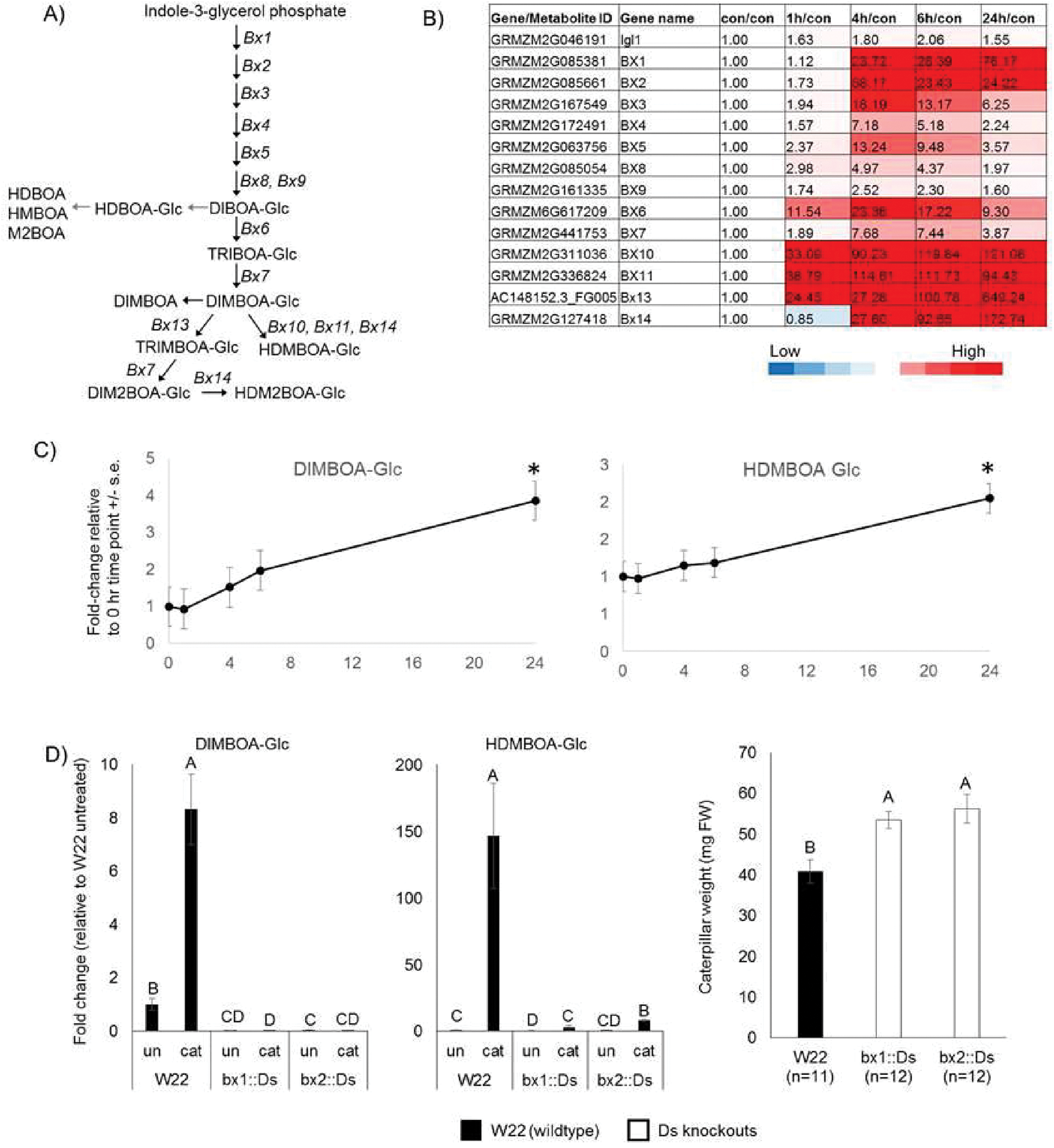
Effects of Caterpillar feeding on benzoxazinoid-related genes and metabolites. A) The benzoxazinoid biosynthesis pathway in maize. B) Heat map of gene expression and C) DIMBOA-Glc and HDMBOA-Glc abundance over time after Caterpillar feeding. Values are presented as fold change relative to untreated control. Mean +/- SE of n = 4 for transcriptomic data, n = 5 metabolic data). D) *S. exigua* Caterpillar body weight after four days on wildtype W22, *bx1::Ds*, and *bx2::Ds* mutant plants. E) Abundance of DIMBOA-Glc and HDMBOA-Glc in wildtype W22, *bx1::Ds*, and *bx2::Ds,* with and without Caterpillar feeding. Different letters above the bars indicate significant differences, *P* < 0.05, ANOVA followed by Tukey’s HSD test.

## CONCLUSION

In this study, we examined the dynamic effects of caterpillar feeding on maize, one of the world’s most important crop plants. Transcriptional and metabolic changes showed rapid responses occurring in the first hour after caterpillar infestation. The transcriptomic and metabolomics changes continue increasing up to the 24 hour time point. Our analysis of the transcriptomic and metabolomic data has led to the characterization of the role of genes from two major defense-related pathways, benzoxazinoids, and jasmonic acid. Future research on these benzoxazinoids and phytohormones that are induced by *S. exigua* feeding in our experiments will enable the breeding of maize cultivars with enhanced resistance to lepidopteran herbivores. In addition, this high-throughput dataset can be utilized to discover key genes that play a role in maize metabolic processes during biotic stress responses.

## SUPPLEMENTARY DATA

**Supplemental Table S1.** Primers used for quantitative RT-PCR analysis.

**Supplemental Table S2.** Primers used to screen for knockout mutations.

**Supplemental Table S3.** RNAseq raw data

**Supplemental Table S4.** RNAseq raw data after data filtering (genes that had expression values of zero more than three times were excluded).

**Supplemental Table S5.** RNAseq data for four caterpillar feeding time points after Cuffdiff.

**Supplemental Table S6.** LC-TOF-MS data from negative ion mode.

**Supplemental Table S7.** LC-TOF-MS data from positive ion mode.

**Supplemental Table S8.** List of differentially expressed maize genes for at least one time point (+/- > 2 fold changed) with *P* value < 0.05, FDR adjusted, used for PLD-SA analysis, clustering over-representation, and PageMan analysis.

**Supplemental Table S9.** Orthologous Arabidopsis and maize genes used for Hormonometer analysis.

**Supplemental Table S10.** RNAseq data of benzoxazinoid genes for four caterpillar feeding time points after Cuffdiff (V3.20).

**Supplemental Table S11.** Bx parameters using LC-TOF-MS.

**Supplemental Figure S1.** Design of the caterpillar feeding experiments. The second leaf of two-week-old B73 maize plants was enclosed in a clip cage. At staggered intervals, two 2^nd^ to 3^rd^ instar *S. exigua* caterpillars were added to each cage. Leaf tissue was harvested after 1 to 24 h of caterpillar feeding. All samples, including controls (untreated), were harvested within a 45-minute time period. Harvested tissue was used for assays of gene expression by Illumina sequencing, as well as for metabolite profiling by LC/MS, and HPLC.

**Supplemental Figure S2.** Comparison of RNAseq and qRT-PCR gene expression data from two independent sets of experimental samples. The following genes were measured by qRT-PCR and RNAseq and normalized to adenine phosphate transferase 1 (*APT1*): GRMZM2G131907, *Bx6* - GRMZM6G617209, Phe Ammonia-Lyase (*PAL*) - GRMZM2G063917, 12-oxo-phytodienoic acid reductase 7 (*OPR7*) - GRMZM2G148281, Lipoxygenase 10 (*LOX10*) - GRMZM2G015419, Lipoxygenase 3 (*LOX3*) - GRMZM2G109130, and Lipoxygenase 8 (*LOX8*) - GRMZM2G104843. Mean +/- SE of n = 4, for RNAseq (orange) and n =5 for qRT-PCR (blue). *R* value = correlation coefficient.

**Supplemental Figure S3.** Venn diagram describing the number of genes up- or down-regulated by caterpillar infestation in 1hr and 24 hr relative to control and 1hr relative to 24 hr. *P* value <0.05 FDR and fold change > 2 or < 0.5.

**Supplemental Figure S4.** Effects of caterpillar feeding on jasmonic acid biosynthesis. A) The jasmonic acid biosynthesis pathway in maize. B) JA and JA conjugate levels of *lox8/ts1-ref* (LOX8; GRMZM2G104843) gene knockout in response to caterpillar attack. Black bars – untreated, white bars- caterpillar infestation for 24 hr. *P < 0.05, two-tailed Student’s *t-*test. ND = not detected.

## ACKNOWLEDGEMENTS

We thank Navid Movahed and Meena Haribal for the technical support with this project. This research was funded by US National Science Foundation awards 1139329 and 1339237 to GJ, and Vaadia-BARD Postdoctoral Fellowship Award FI-471-2012 to VT.

